# RPRH: a web-based tool for the inference of potential roles of plant microRNAs in human body

**DOI:** 10.1101/2020.06.11.145367

**Authors:** Yuanxu Gao, Chunmei Cui, Kaiwen Jia, Yuan Zhou, Qinghua Cui

## Abstract

Recently, increasing studies reported that plant microRNAs (miRNAs) from food could be absorbed through gastrointestinal tract to regulate the physiological processes of animals. These novel cross-kingdom regulatory roles of plant miRNAs open a new field in miRNA study and drug discovery, but the lack of corresponding bioinformatics tools limit the development of this field. In this study, we built a web-based tool, RPRH (Role of Plant microRNA in Human), to infer the potential roles of plant miRNAs in human and to profile the landscape of all documented plant miRNAs regulatory functions in human. Totally, RPRH included 10414 miRNAs from 82 plant species and 18062 annotation terms like diseases, pathway, gene ontology (GO), and drugs. RPRH represents a bioinformatics resource for further investigation of cross-kingdom regulations of plant miRNAs.

**Availability:** http://www.rnanut.net/rprh/

## Introduction

MicroRNAs (miRNAs) are a class of small noncoding RNAs involved in the post-transcriptional regulation of gene expression by targeting 3’UTR of mRNA. These molecules play important roles in various biological processes including but not limited to cell proliferation, differentiation, and apoptosis (*1*). miRNAs could not only target endogenous genes in cells, but also exist stably in many body fluids such as blood circulation (*2*). Recently, increasing evidence revealed that plant miRNAs could be absorbed into mammalian through food intake to regulate physiological and pathological process in hosts, indicating that plant miRNAs hold great promise for future biomedical research (*3–5*). Such phenomena are highlighted as the cross-kingdom regulation of plant miRNAs (*3*). It is believed that these plant-derived miRNAs have roles in human biological processes and disease (*6*). In 2007, we built the human miRNA disease database (HMDD) (*7*) and proposed version 2 and 3 in 2014 and this year (*8, 9*). HMDD collected the associations between human disease and human miRNAs but did not collect plant miRNAs involved in human disease. However, dissecting the roles of plant miRNAs in human body is important. To date, no systematic bioinformatics study on the cross-kingdom regulation between plant miRNAs and human is available.

There are a few bioinformatics works focus on the miRNA cross-kingdom regulation. Pirrò (*10*) developed a tool named MirCompare to infer plant miRNAs which have the same regulatory functions of mammalian endogenous miRNAs based on the similarity of overall sequences and seed regions between miRNA pairs. However, it is hard to find out potential roles of plant miRNAs who do not have similar endogenous miRNAs. In addition, Zhang et al. presented a method for constructing regulatory networks according to predicted target genes both in plant and human to study cross-kingdom regulation, and applied their method to miRNAs from *Arabidopsis thaliana* (*11*). Nonetheless, plants, especially agricultural species like crops and vegetables, have a wide spectrum of miRNAs encoded in their genomes, but there is no research profiling the landscape of the roles of plant miRNAs in human. For this purpose, here we first proposed a computational method and then built a database for dissecting the roles of plant miRNAs in human body.

## Materials and methods

### Data collection and curation

Mature miRNA sequence data were downloaded from miRBase (release 22) (http://www.mirbase.org). Plant miRNAs, as of whom the species belong to Viridiplantae in the National Center for Biotechnology Information taxonomy database, were selected by manual screening. Human 3’UTR sequences were obtained from the UCSC genome browser (http://genome.ucsc.edu/cgi-bin/hgTables, genome version GRCh38) and the TargetScan website (http://www.targetscan.org/vert_72/vert_72_data_download/UTR_Sequences.txt.zip). The 3’UTR sequences getting from UCSC were filtered and only those located in coding genes were kept. Annotated functional gene sets (diseases, GO, pathway etc) were download from GeneSetDB (*12*), MsigDB (*13*) and collected from literature (*14*). We organized all gene sets and divided them into five subclasses, which include Pathway, Disease, Drug, Phenotype, and Gene Ontology. Experimental validated human microRNA targets were download from miRTarBase (http://mirtarbase.mbc.nctu.edu.tw/cache/download/7.0/hsa_MTI.xlsx) (*15*).

### Plant miRNA target prediction

The target genes of plant miRNAs in human were predicted separately by three tools, TargetScan (*16*), miRanda (*17*), and PITA (*18*). Prediction by miRanda used UCSC 3’UTR as UTR input with default settings. Prediction by PITA used UCSC 3’UTR as UTR input and in order to improve prediction accuracy, we optimized the parameter as “−l 7-8 -gu ‘7;18;1’ -m ‘7;0,8;0’” and set cutoff of the final score as −10. As the prediction accuracy is better when using the provided 3’UTR sequences which include cross-species conservation information, prediction by TargetScan used TargetScan 3’UTR file as UTR input with default settings. All the prediction targets were convert transcripts ID into gene symbol. And the target gene set for each miRNA was defined as the consensus between three prediction tools.

### Gene set enrichment analysis

We performed gene set enrichment analysis with the organized gene sets (Pathway, Disease, Drug, Phenotype, and Gene Ontology) by using the hypergeometric test. Assuming that *A* represents the number of all coding genes, *S* represents the number of genes in signature gene set *x*, *T* represents the number of target genes of given miRNAs and *TS* represents the number of miRNA target genes included in *S*. Statistical significance of gene set was calculated as the following formula.

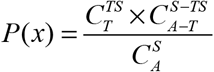

The symbol “*C*” means the combination operation. Finally, the P values for all signature gene sets are adjusted by FDR corrections.

### Endogenous miRNAs similarity calculation

The similarity between plant miRNA and human endogenous miRNA were characterized using Jaccard similarity between their targets. Assuming that *A* represents the plant miRNA target gene sets and *B* represents the human endogenous miRNA target gene sets. The similarity was calculated as the following formula.

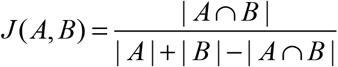

We normalized the Jaccard index among all possible plant miRNA and human miRNA pairs by using Z-score method. The significance threshold was set as 1.96, which is corresponding to P < 0.05.

## Results

### The framework for dissecting the roles of plant miRNAs in human body

The workflow includes data collection, target prediction, results integration and function analysis (Fig. 1). Firstly, we collected and sorted plant miRNA sequences, human 3’UTR sequences and functional gene sets. Then we predicted the targets of each miRNA using three prediction tools. Next, we convert the transcript ID into gene symbol and take the consensus of results between three tools as target gene set. Finally, we perform gene set enrichment analysis with organized functional gene sets to figure out potential roles of plant miRNA.

**Figure 1.**
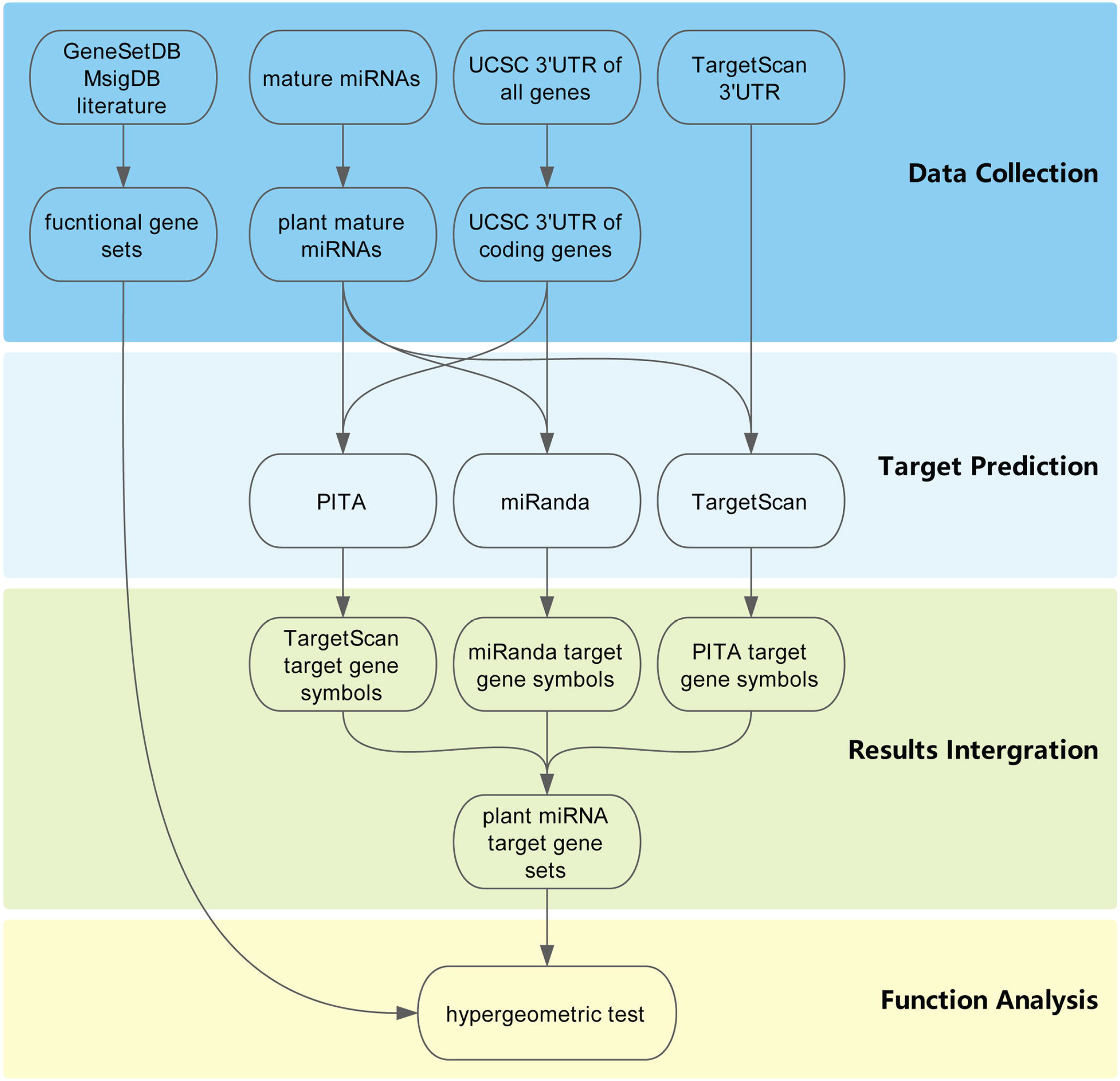
The framework for predicting potential roles of plant miRNAs in human.

### Collection of Plant miRNAs and human 3’UTR sequences

To infer the potential roles of plant miRNAs in human bodies, we began by collecting and sorting plant miRNA data and human 3’UTR data. As a result, the plant miRNA data consist of 10414 mature plant miRNA sequences across 82 plant species (Fig. 2A). The top fifteen species and families with the most miRNAs are depicted in Fig. 2B and 2C, of which most species are model organism and well-studied crops. 65 of 82 plant species have less than 200 recorded miRNAs including many crops and vegetables like *Triticum aestivum*, which suggests a scarcity of the microRNA study on these species. The lengths of mature plant miRNAs range from 17nt (nucleotides) to 28nt (Fig. 2D) and the longest miRNA is cst-miR11332 from *Cucumis sativus.* Most plant miRNAs (6428/10414) are 21 nucleotides in length which is consistent with previous observations. For 3’UTR sequence data, we obtained 42364 3’UTR sequences from UCSC and 28352 3’UTR sequences from TargetScan.

**Figure 2.**
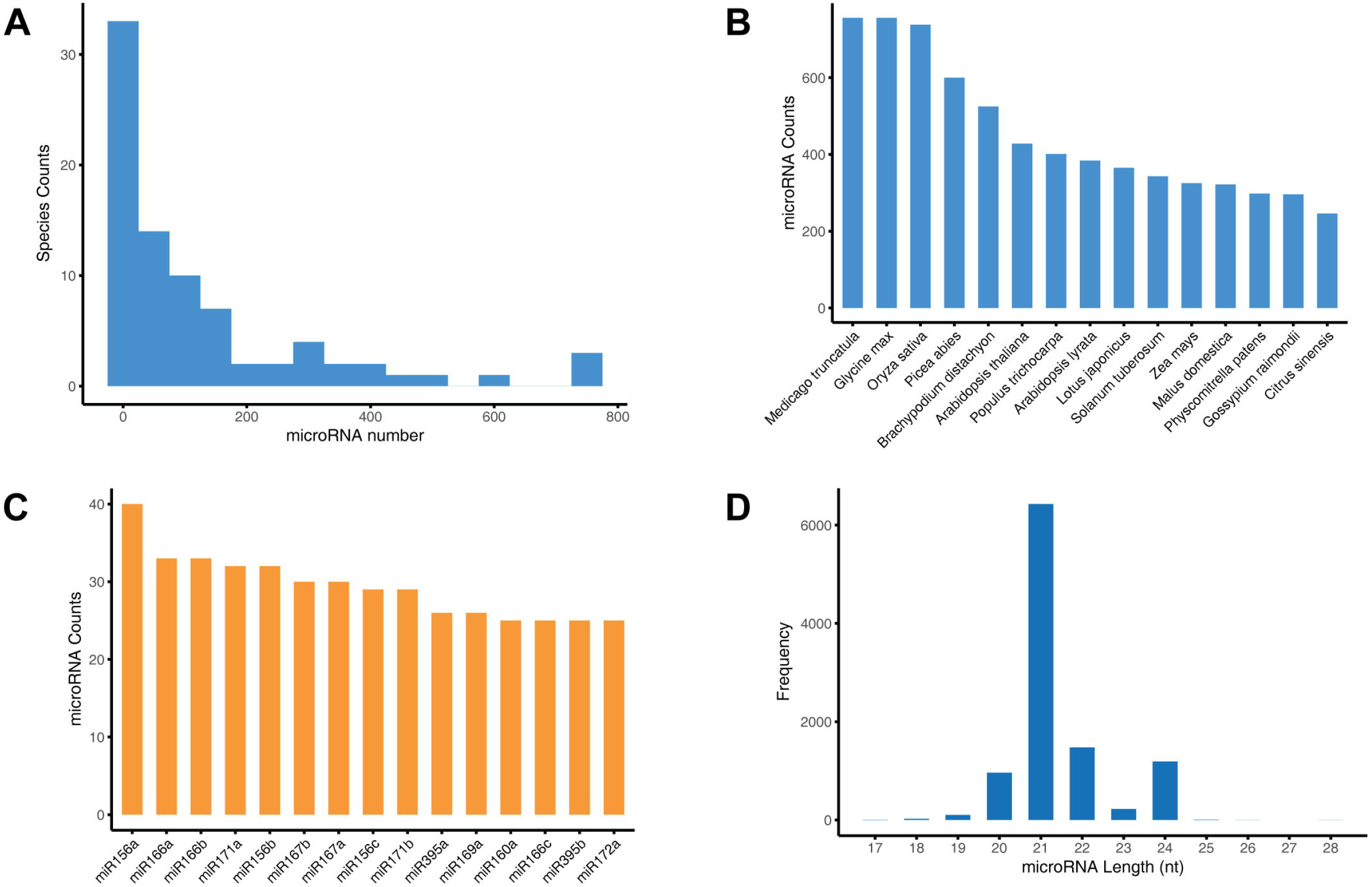
Statistics of plant miRNA data. (A) Distribution of miRNA number of plant species. (B) The top 15 species with the largest number of miRNAs. (C) The top 15 families with the largest number of miRNAs. (D) Distribution of miRNA lengths.

### Prediction of plant miRNAs target genes in human

Then we try to predict the target genes of plant miRNAs in human. According to the experimental evidence (*3*), plant miRNAs play post-transcriptional regulatory role in a fashion mimicking the endogenous miRNAs in mammalian. We performed target prediction using three software including TargetScan (*16*), miRanda (*17*) and PITA (*18*), separately. For each plant miRNA, we take the consensus between three prediction results as the finally target gene set. The distribution of target gene number of plant miRNA is shown in Fig. 3A. On average, there are 570 target genes for each plant miRNA. miRNAs with most target genes are ptc-miR7836 and ptc-miR7837 which targeting 4535 genes. On the contrary, miRNAs with fewest target genes are nta-miR6148a and eun-miR10214a-5p which targeting only 1 gene. Prediction results of corresponding software are also shown in Fig. 3B-D.

**Figure 3.**
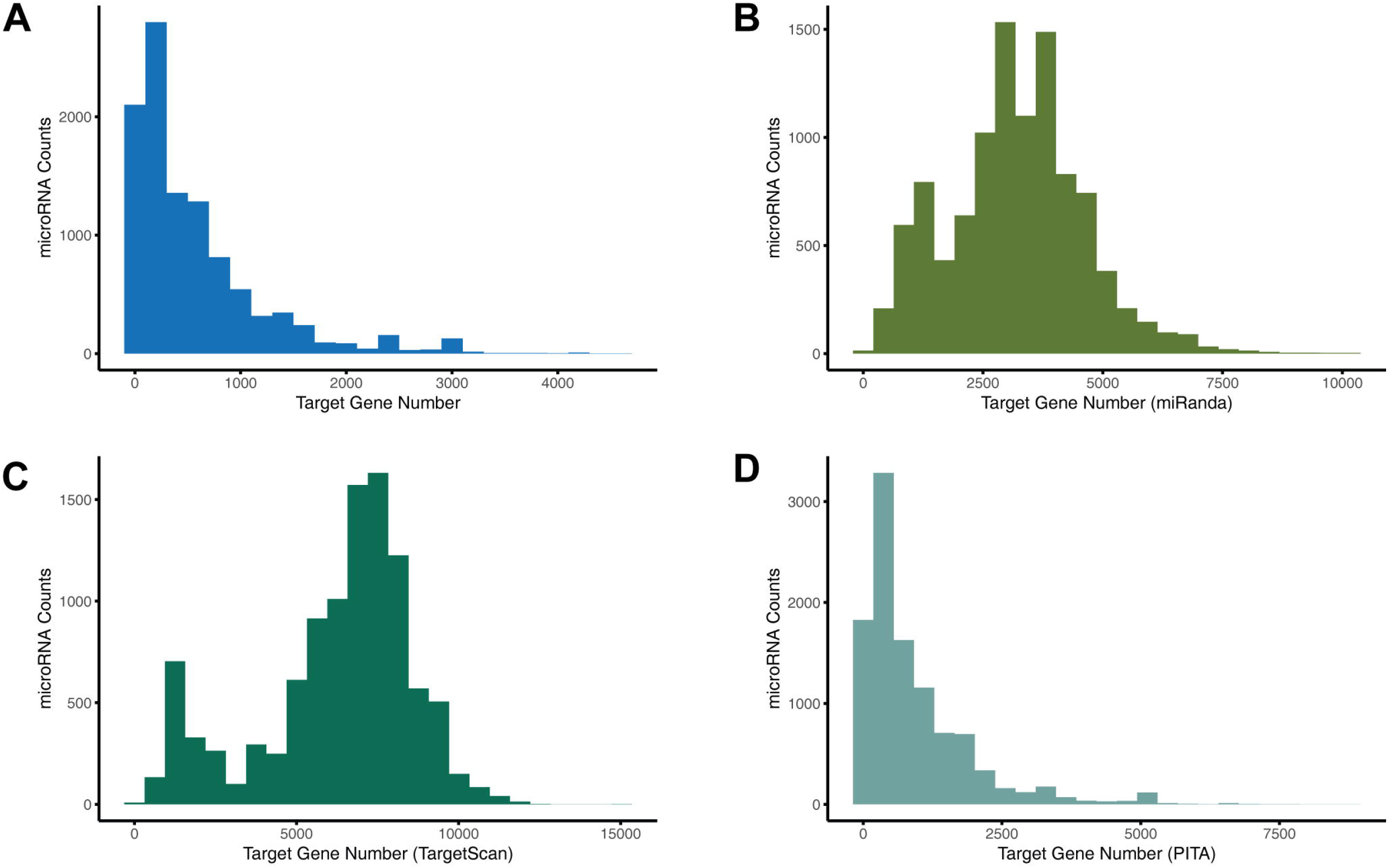
Statistics of predicted plant miRNA-human target. (A) Distribution of the number of target genes of plant miRNAs. (B) Distribution of the number of miRanda target genes. (C) Distribution of the number of TargetScan target genes. (D) Distribution of the number of PITA target genes.

### Inference of potential roles of plant miRNAs in human bodies

In order to obtain a comprehensive overview of the potential roles of plant miRNAs, we organized an integrated database of functional gene sets which contain data obtained from GeneSetDB (*12*), MsigDB (*13*) and literature (*14*). All the gene sets were divided into 5 subclass consisting of pathway, disease, drug, phenotype and gene ontology based on their source databases (Fig. 4A). Then we perform function enrichment analysis on target gene set of each plant miRNA with the organized functional gene sets. As a result, we obtained 40743425 miRNA-functional annotation associations. Most miRNAs (86.2%) have less than 5000 annotations (Fig. 4B). We noted that our method could capture the known cross-kingdom regulations of plant miRNAs. For example, osa-miR168a-5p was experimental proved that it could regulate glycolipid metabolism in mammals (*3*). In our study, osa-miR168a-5p was found involved in positive regulation of glucose metabolic process in human (P = 5.15e-4). Another research showed that plant miR159 could participate in WNT signal pathway and suppresses proliferation of breast cancer cells (*5*). In our study, soybean (*Glycine max*) microRNA gma-miR159a-3p were significant enriched for WNT signaling pathway (P = 4.28e-3) in pathway subclass and breast secretory ductal carcinoma (P = 3.26e-2) in diseases subclass. As plant miRNA perform their regulatory roles in a way similar to endogenous miRNA, there might be human miRNAs that target the same gene, which make plant miRNA likely have no action (Fig. 4C). To address this issue, we calculated the similarity between plant miRNA and human endogenous miRNA based on their target gene. A total of 2390 plant miRNAs have no similar endogenous miRNA, which indicating that they may perform a different role in human body (Fig. 4D).

**Figure 4.**
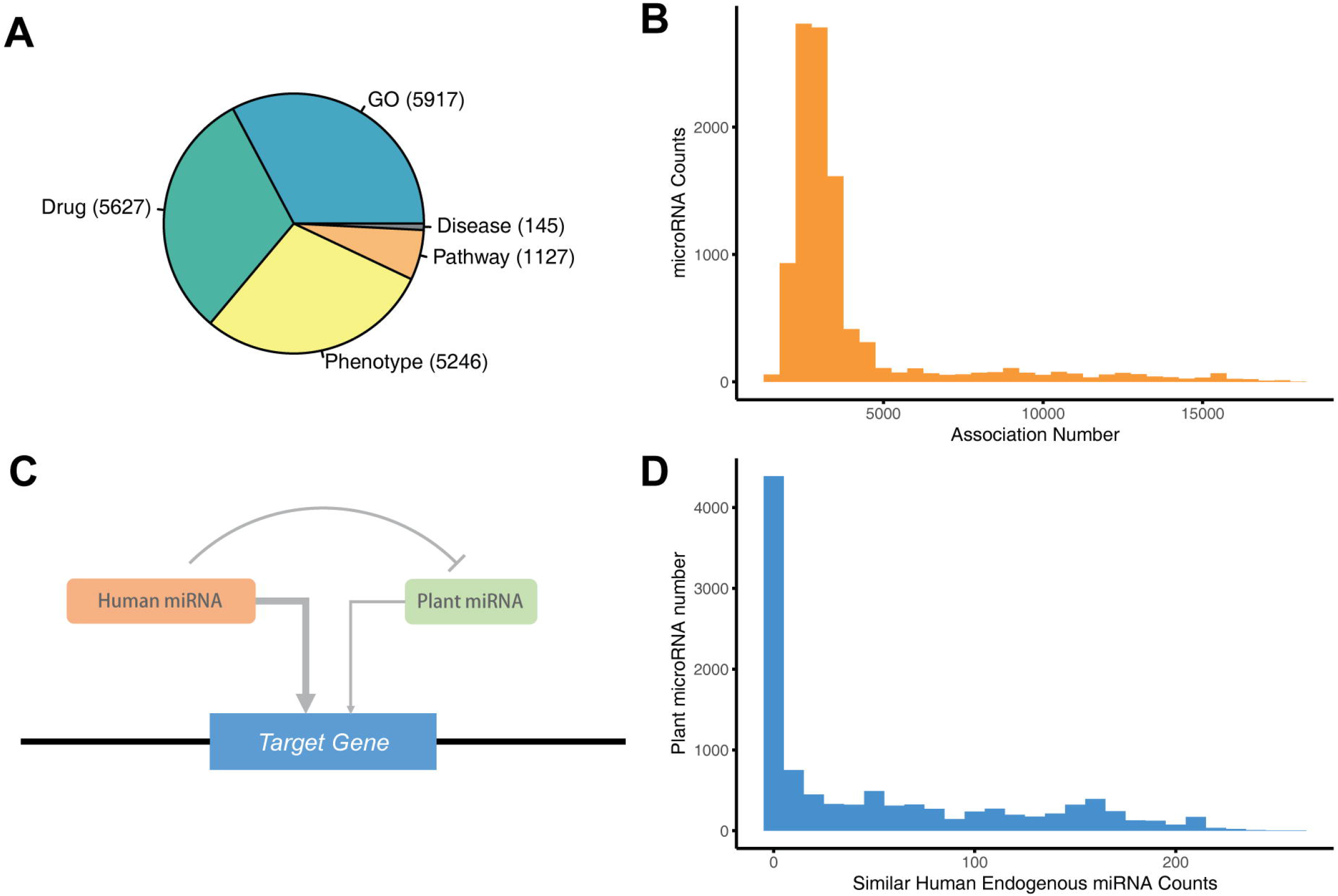
Statistics of predicted gene set terms of plant miRNAs. (A) Statistics of gene set term subclasses. (B) Distribution of the number of predicted gene set associations of plant miRNAs. (C) Schematic diagram of relationship between plant miRNA and similar human endogenous miRNA. (D) Distribution of the number of human endogenous miRNAs similar to plant miRNA.

### The construction of the RPRH web server

To facilitate users to use our method, we built the RPRH web server which is available at http://www.rnanut.net/rprh. All prediction results were organized by MariaDB, an open source database management system. And the website was developed based on CodeIgniter, a PHP web framework. Users could search the database by plant miRNA names to see their potential roles in human body. Our server supports fuzzy search and is case-insensitive for input. After searching a specific plant miRNA, a search result page will return to show all documented plant miRNAs that are similar to input for users to choose. In result page, all gene set terms are shown by subclass and ordered by significance of enrichment analysis (Fig. 5A). We also provide the download options for users so they could download the target genes and all enriched gene set terms of the given plant miRNA. Similar human endogenous miRNAs of each plant miRNA are shown in a standalone page and are organized in a network where the center node represents plant miRNA, peripheral nodes represents human miRNAs and the thickness of lines represents the target similarity between two miRNAs (Fig. 5B). Besides, if users are interested in human genes and try to find out potential regulatory plant miRNAs, they could search by a human gene symbol which also supports fuzzy search. If there are no results for a candidate plant miRNA, website will have a predict page for users to predict the targets of their own miRNA. Moreover, the target genes of all plant miRNAs in our database can be download at the “Download” page. In addition, a tutorial for the usage of the tool is available in the “Help” page.

**Figure 5.**
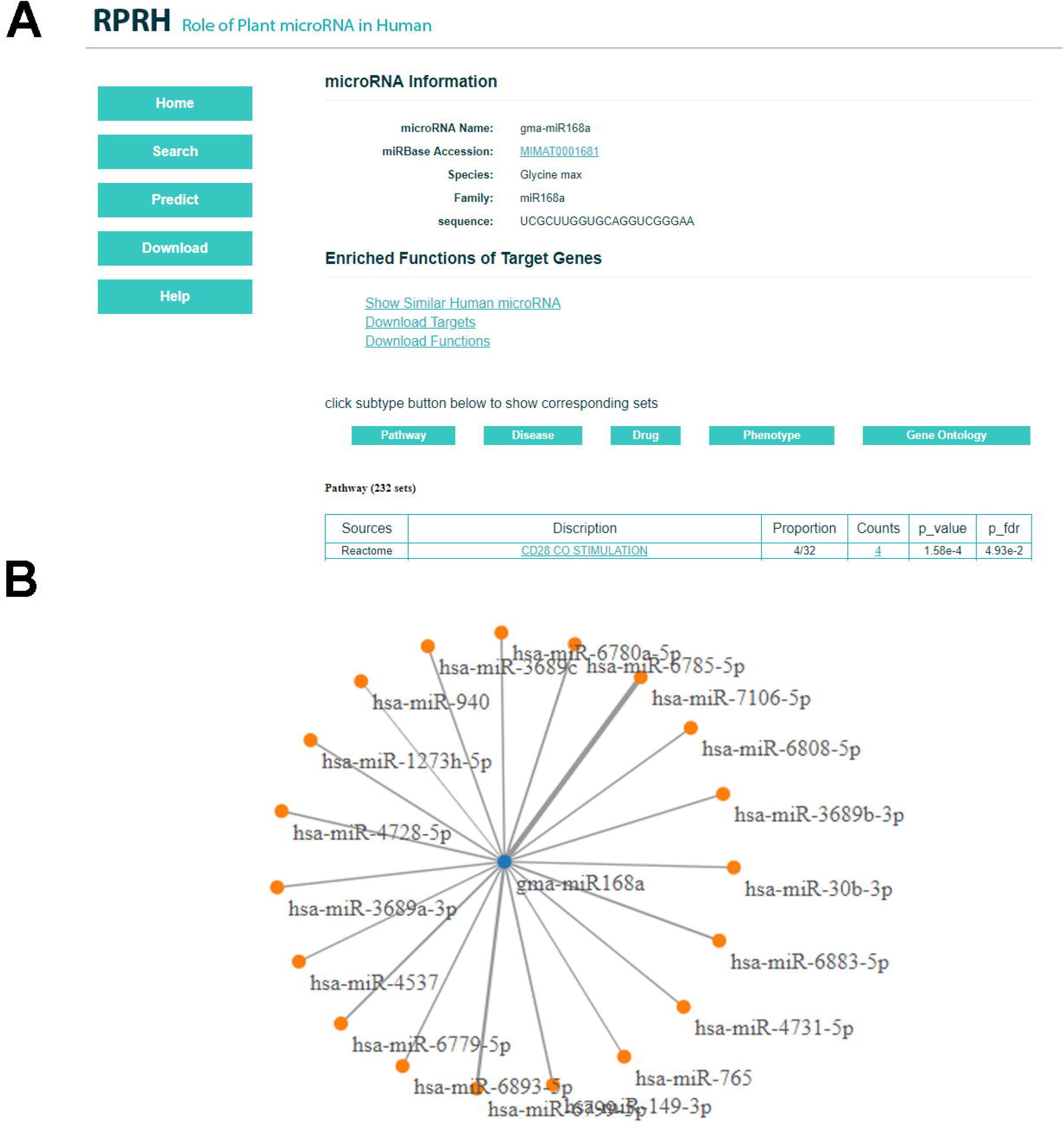
Screenshots of the RPRH web server. (A) Search result page. (B) Network of plant miRNA and similar human endogenous miRNAs.

## Discussion

The inter-species regulation of miRNAs has already been shown in many virus and parasites (*6*). Early study in 2004 shown that virus could encode miRNAs and regulate host gene expression (*19*). Recently research found that many 22 nucleotides miRNAs from *Cuscuta campestris*, a parasitic plant, could act as trans-species regulators to affect the host gene expression (*20*). However, interaction between plant miRNAs and mammalian is novel and attractive for its potential in medicine(*21*). Here we built the web-based tool, RPRH, to comprehensively profile the potential roles of plant miRNAs in human body. We note that it remains largely unclear how plant miRNAs regulate human genes. Here, based on the current knowledge, we use the tools for mammalian miRNA target prediction to identify the human target genes of plant miRNAs. If special rules and mechanisms for cross-species miRNA-target regulation are found in the future, novel algorithms should be developed for such special cases. Nevertheless, although limitation exists in the current stage, we believe that the RPRH represents a valuable resource to investigate the novel cross-kingdom regulations of miRNAs.

